# Hypothalamic vasopressin sex differentiation is established during embryonic neurogenesis

**DOI:** 10.1101/2022.11.17.516788

**Authors:** Jing Zheng, Parisa Moazen, Dinara Baimoukhametova, Catherine Lebel, Jaideep Bains, Deborah Kurrasch

## Abstract

Arginine vasopressin (AVP) is a conserved sexually dimorphic system regulating behaviors and physiologies. Decades of research concludes that AVP sex differentiation in mice occurs around birth when a gonadal surge of testosterone is converted to estrogen in male sexual dimorphic brain regions. We discovered that AVP neurons in the murine hypothalamus are sexually dimorphic much earlier than this surge, with male mice displaying more AVP neurons than females during neurogenesis at E11. *In utero* exposure to a pan-ER antagonist blocked AVP masculinization in males, whereas embryonic exogenous 17β-estradiol or xenoestrogen bisphenol A (BPA) masculinized female AVP neurons, causing permanent masculinization effects on adult female AVP cell numbers, projections and intrinsic neuronal properties. Our data reveal that sexual dimorphism of hypothalamic AVP neurons is present from the earliest stages of their differentiation and challenges current dogma that masculinization of the male brain occurs around birth.

**One-Sentence Summary:** Sexual dimorphism of hypothalamic vasopressin is established during neurogenesis and exposure to the xenoestrogen bisphenol A during pregnancy induces masculinization of female brains

## Main Text

Arginine vasopressin (AVP) circuits are evolutionarily conserved and play a role in social behaviors such as aggression, parenting, and play, as well as homeostatic processes such as thirst and fluid retention (*1*). The AVP system is one of the most consistently sexually dimorphic neural systems across taxa, with *Avp* transcripts, neuronal numbers, and projection densities higher in males than in females in all species studied except fish (*2*). AVP neurons and/or circuits are sexually dimorphic across several brain regions including the lateral septum (LS), the bed nucleus of stria terminalis (BNST), medial amygdala (MeA), and lateral habenula (LH) (*3, 4*). Within the hypothalamus, AVP neurons are localized in the paraventricular nucleus (PVN), suprachiasmatic nucleus (SCN), and supraoptic nucleus (SON) (*5*), although the extent of sex dimorphism in these regions is debated (*6, 7*). In humans, disruptions of AVP circuits are widely associated with neurological (*8, 9*) and neurodevelopmental (*10*) disorders.

Our current understanding of neural sex differentiation arises from classic experiments in rodents conducted over fifty years ago whereby testes transplanted into the necks of genetic females at birth induced masculine behavioral patterns and castration of genetic males at birth resulted in feminine behaviors (*11*). Decades of studies across various species have entrenched the idea that neural sex differentiation is driven by a testicular testosterone surge around birth, with testosterone converted to estradiol by aromatase-expressing cells in sexually dimorphic brain regions in males (*12–15*), thereby counter-intuitively making estrogen masculinizing during this critical window. Thus, according to doctrine, the timing for sexually dimorphic programs is the perinatal period (~E17-P2) when the gonads have fully differentiated to produce physiological levels of circulating testosterone (*16*).

Here we studied the timing of sex differentiation of AVP neurons within the PVN region of the embryonic hypothalamus (hereafter AVP^PVN^), using 17β-estradiol and bisphenol A (BPA) to modulate estrogen signaling. BPA is a plasticizer widely used in the production of consumer products (*17*) and disrupts endocrine systems by acting as a weak estrogen agonist (*18, 19*).

## Results

### Sex dimorphism of AVP^PVN^ neurons is established during neurogenesis and requires estrogen

In the embryonic brain, we observed significantly higher numbers of AVP+ cells in the male versus female PVN at E14.5 (Fig. 1A-B), days prior to any gonadal testosterone surge. *In utero* injection at E13.5 of the pan-ER antagonist fulvestrant (4 μM, 16 μM) into the adjacent 3^rd^ ventricle dose dependently prevented masculinization, with males acquiring female-like numbers of AVP+ neurons (E15.5, Fig. 1C-D). *In utero* injection of 17β-estradiol (E14.5-E15.5; E2, 4 μM) or gestational exposure to the weak estrogen BPA (E0.5-E15.5; 2.25 μg/kg bw/day as per (*20*)) masculinized the female brain, with females displaying male-like numbers of AVP+ cells compared to female control (Fig. 1E-F). Curiously, exogenous exposure to estrogen or gestational BPA (E0.5-E15.5) in males caused a decrease in AVP^PVN^ neuronal numbers (Fig. 1E-F). Restricting BPA exposure to the neurogenic window (E7.5-E15.5) caused male-like numbers of AVP+ neurons in the perinatal and juvenile female brain, as observed above, and no sustained masculinization of the male brain (Fig. 1G-H). We employed gestational BPA at environmentally relevant levels as an estrogenic tool for the remainder of the studies.

**Fig. 1.**
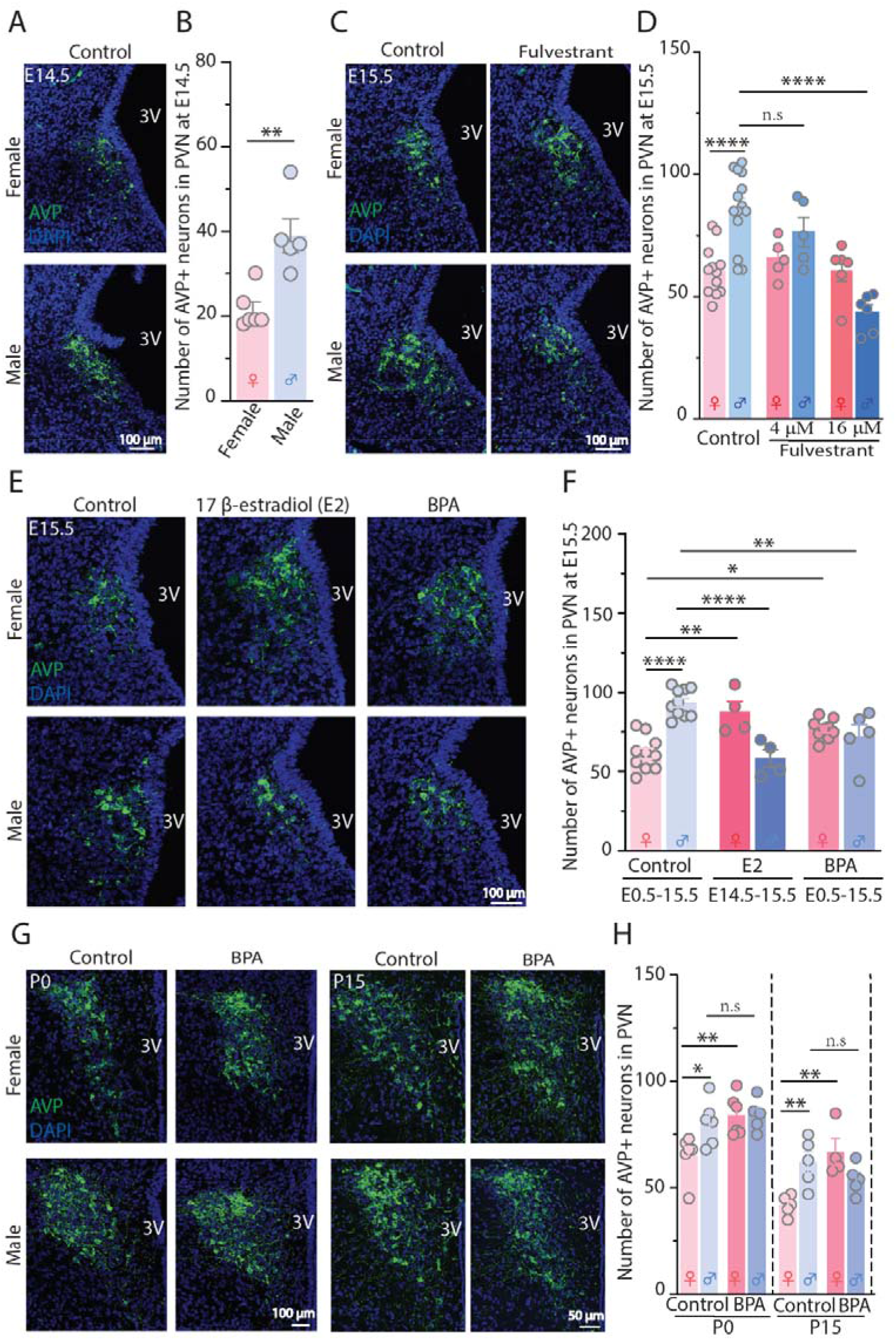
Sexual dimorphism of AVP^PVN^ initiates in the embryo. (**A** to **B**) Representative images and number of AVP+ neurons in the AVP^PVN^ of female and male embryonic (E) 14.5 brain; n = 5-6 mice from 2 litters, with two-side unpaired t- test. (**C** to **D**) The impact of embryonic exposure (E13.5) to corn oil (Control) or the pan-ER antagonist fulvestrant (4 μM and 16 μM) on the number of AVP^PVN^ neurons in female and male brains at E15.5; n = 5-14 mice from 2-3 litters. The representative images (**E**) and number (**F**) of AVP^PVN^ neurons in female and male brains at E15.5 in the presence of 17β-estradiol (E2, 4 μM; in utero injection at E14.5) or bisphenol A (BPA, 2.25 μg/kg bw/day; maternal exposure from E0.5) embryonic exposures; n = 4-10 mice from 2-5 litters. (**G** to **H**) The effect of maternal BPA exposure (from E7.5 to E15.5) on the number of AVP^PVN^ neurons in female and male brains at postnatal (P) 0 and P15 day; n = 5-6 mice from 2-3 litters. AVP, arginine vasopressin; DAPI, nuclei marker; 3V, 3^rd^ ventricle. *\*P* < 0.05, \*\**P* < 0.01, \*\*\*\**P* < 0.0001, n.s. = no significance; data shown expressed as mean ± SEM; data were analyzed with one-way ANOVA followed by Sidak’s multiple comparison correction, unless otherwise stated.

Next, we conducted bromodeoxyuridine (BrdU) birthdating to test if males displayed higher numbers of AVP+ neurons from neuronal birth. We observed significantly more AVP^PVN^ neurons born in males compared to females at peak neurogenesis (E11), followed by equal numbers of AVP+ neurons between males and females born at E12 and E13 (Fig. 2A-E). Gestational BPA treatment did not alter the number of AVP+ neurons born in exposed males but caused male-like numbers of AVP^PVN^ neurons to be born at E11 and E12 in BPA-exposed females (Fig. 2D-E). These findings suggest that the sexually dimorphic pattern of AVP^PVN^ neurons is established during neurogenesis.

**Fig. 2.**
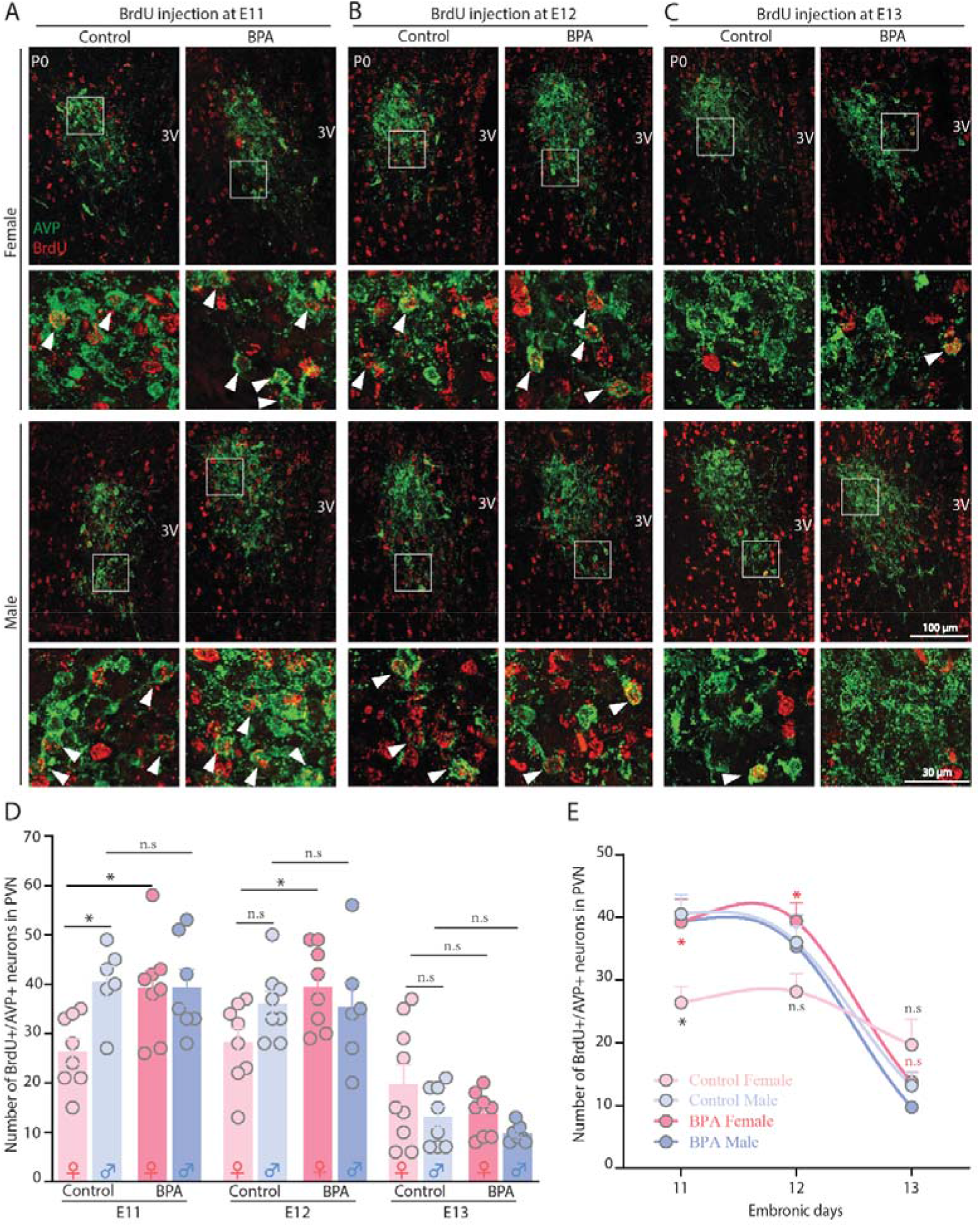
AVP sexual dimorphism is established during neurogenesis and is reversed by maternal exposure to BPA. (**A** to **C**) Representative P0 images of AVP^PVN^ neurons in female and male brains maternally exposed to corn oil (Control) or BPA showing newborn AVP neurons at E11, E12 and E13 visualized by AVP (green) and BrdU (red) colocalization. White squares indicate region of zoom-in images, with arrowheads pointing to the AVP+/BrdU+ neurons. 3V, 3 ventricle. (**D**) Quantitative bar graphs showing the number of AVP^PVN^ neurons born on day of BrdU injection in females and males at E11, E12 and E13, respectively, in the presence of maternal exposures to corn oil (Control) or BPA. n = 6-8 mice from 2 litters/sex/group. (**E**) Fitted curves based on data from (D) showing sexually dimorphic pattern of AVP neurogenesis and the disruption of maternal BPA exposure on this timing. P values in black compare control female versus control male, while the values in red compare control female versus BPA female. *\*P* < 0.05, n.s = no significance; data expressed as mean ± SEM, with all data were analyzed using one-way ANOVA followed by Sidak’s multiple comparison correction, except the E12 data that were analyzed using Brown-Forsythe and Welch ANOVA followed by Tamhone’s T2 multiple comparison.

### Masculinization of female AVP^PVN^ neurons from gestational BPA exposure is maintained in the postnatal, juvenile, and adult stages

We next tested the long-term organizing effect of embryonic sex differentiation by examining whether gestational BPA-induced masculinization of females was maintained after birth. The numbers of AVP^PVN^ neurons were quantified at birth (P0), postnatal (P7), juvenile (P15 and P30), and adult (4 months) stages for males and females exposed to control or gestational BPA (E0.5-P0; 2.25 μg/kg bw/day). In control animals, we observed significantly more AVP^PVN^ neurons in control males than in females across all stages tested (Fig. 3A-D), demonstrating sex dimorphism of hypothalamic PVN vasopressin system, a debated topic. Next, we examined the lasting effects of gestational BPA and observed an increased number of AVP+ neurons in exposed females that equals control male numbers across all time points, consistent with a permanent organizing effect. In contrast, gestational BPA decreased the number AVP+ neurons at birth in males, as observed in Fig. 1D; however, the dimorphic levels were restored by P7, P15, P30 and 4 months (Fig. 3A-D). Together, these data suggest that embryonic masculinization of female AVP^PVN^ neurons results in permanent sex dimorphisms in the adult.

**Fig. 3.**
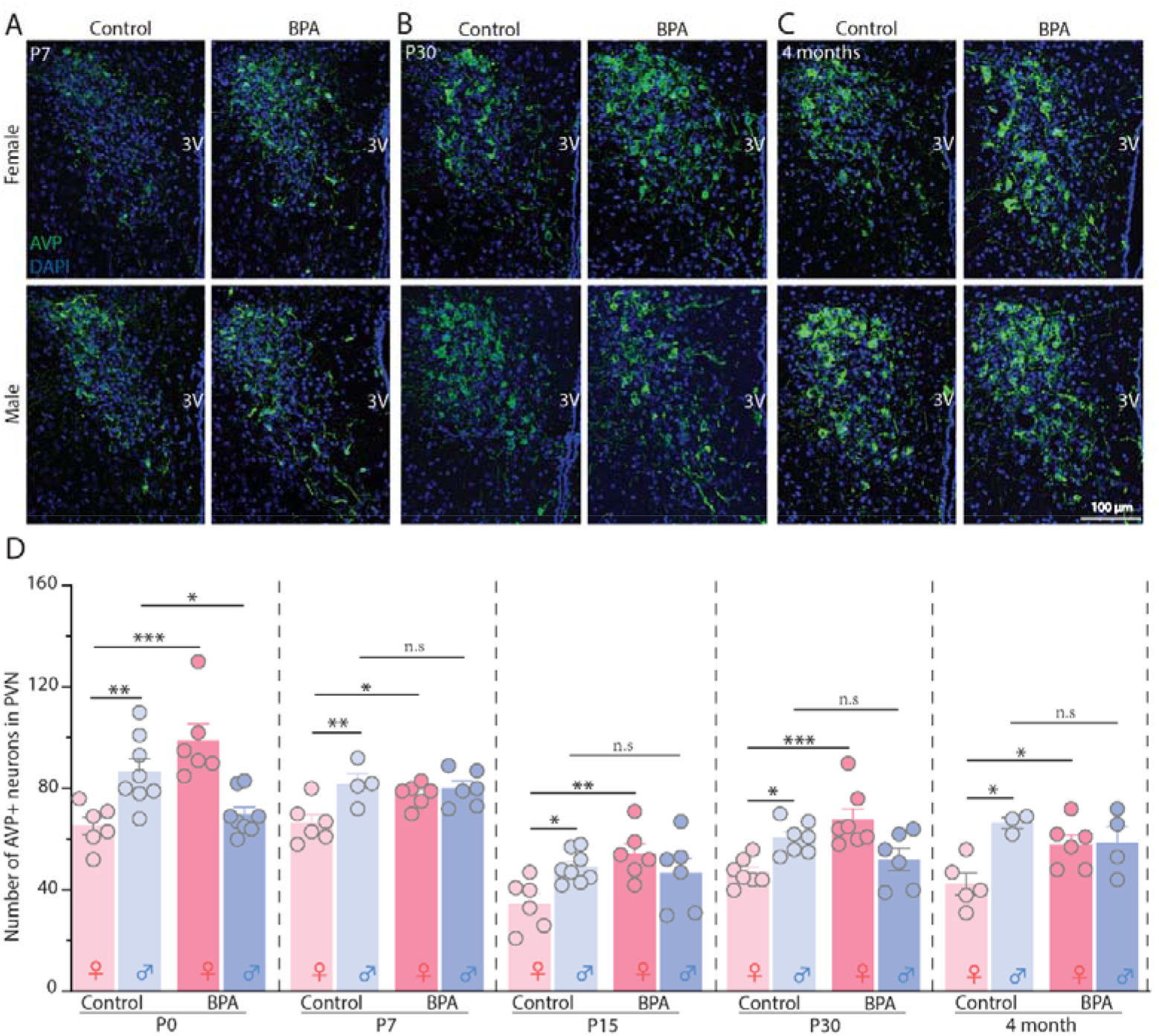
Masculinization of female AVP^PVN^ neurons from gestational BPA exposure is maintained in the postnatal, juvenile, and adult female hypothalamus. (**A** to **C**) Representative images of P7, P30 and 4 month old PVN stained with anti-vasopressin antibody (green) and nuclei marker (DAPI, blue) in female and male maternally exposed to corn oil (Control) or BPA. 3V, 3^rd^ ventricle. (**D**) Quantitative results of the number of AVP+ neurons at each stage in the absence (Control) or presence of BPA exposure in female and male mice. n = 3-8 mice from 2-3 litters/sex/group. \**P* < 0.05, \*\**P* < 0.01, \*\*\**P* < 0.001, n.s. = no significance; data expressed as mean ± SEM, with data analyzed using one-way ANOVA followed by Sidak’s multiple comparison correction.

To determine if this sex dimorphism was consistent across hypothalamic vasopressin populations outside the PVN, we also quantified AVP+ neurons in the E15.5 and P30 hypothalamic SON. We observed sex dimorphism in AVP neurons in control male versus females in the P30 but not E15.5 SON (fig. S1A-D). Gestational BPA exposure increased AVP+ neurons in both males and females at P30 but only in female SON at E15.5 (fig. S1A-D). Additionally, we examined if other hypothalamic PVN cell types were sex dimorphic, namely corticotropin-releasing hormone (CRH)+ neurons and AVP+/CRH+ dual-expressing cells at P7 and P30. We did not observe a significant difference in the number of CRH+ neurons between male and female in the P7 and P30 PVN, and gestational BPA exposure elevated CRH+ cells in only P30 males (fig. S1E-H). No differences in the number of AVP+/CRH+ neurons in the PVN of male and female control brains were observed in P7 and P30 mice; however, gestational BPA exposure caused a substantial decrease in both P30 male and female AVP+/CRH+ cells compared to control (fig. S1I-L), suggesting that BPA-induced masculinization of female AVP+ cells might arise at the expense of AVP+/CRH+.

### Gestational BPA exposures disrupts sexual dimorphism of AVP projections in intra- and extra-hypothalamic regions

AVP^PVN^ neurons regulate social behaviors by targeting intra-hypothalamic projections to the SON, as well as extra-hypothalamic projections to regions such as the septal area, BNST, MeA and LH. We observed an increase in intra-hypothalamic AVP- immunoreactivity (AVP-ir) between the PVN and SON in control males compared to females (Fig. 4A-B). Gestational BPA exposure caused masculinization of female PVN AVP-ir to the SON as well as a decrease in male AVP-ir (Fig. 4A-B), concomitant with the increase or decrease in AVP+ cell numbers observed above, respectively (Fig. 3A-D). We confirmed these findings using whole brain iDISCO + 3D imaging, where an increase in male AVP-ir between the PVN and SON and masculinization of female projections was clearer (Fig. 4C, asterisks). iDISCO+ imaging also revealed that the dorsomedial hypothalamus might be an unappreciated sexually dimorphic region, where males displayed denser AVP projections than females fig. S2D). This pattern was disrupted by maternal BPA exposure (fig. S2D).

**Fig. 4.**
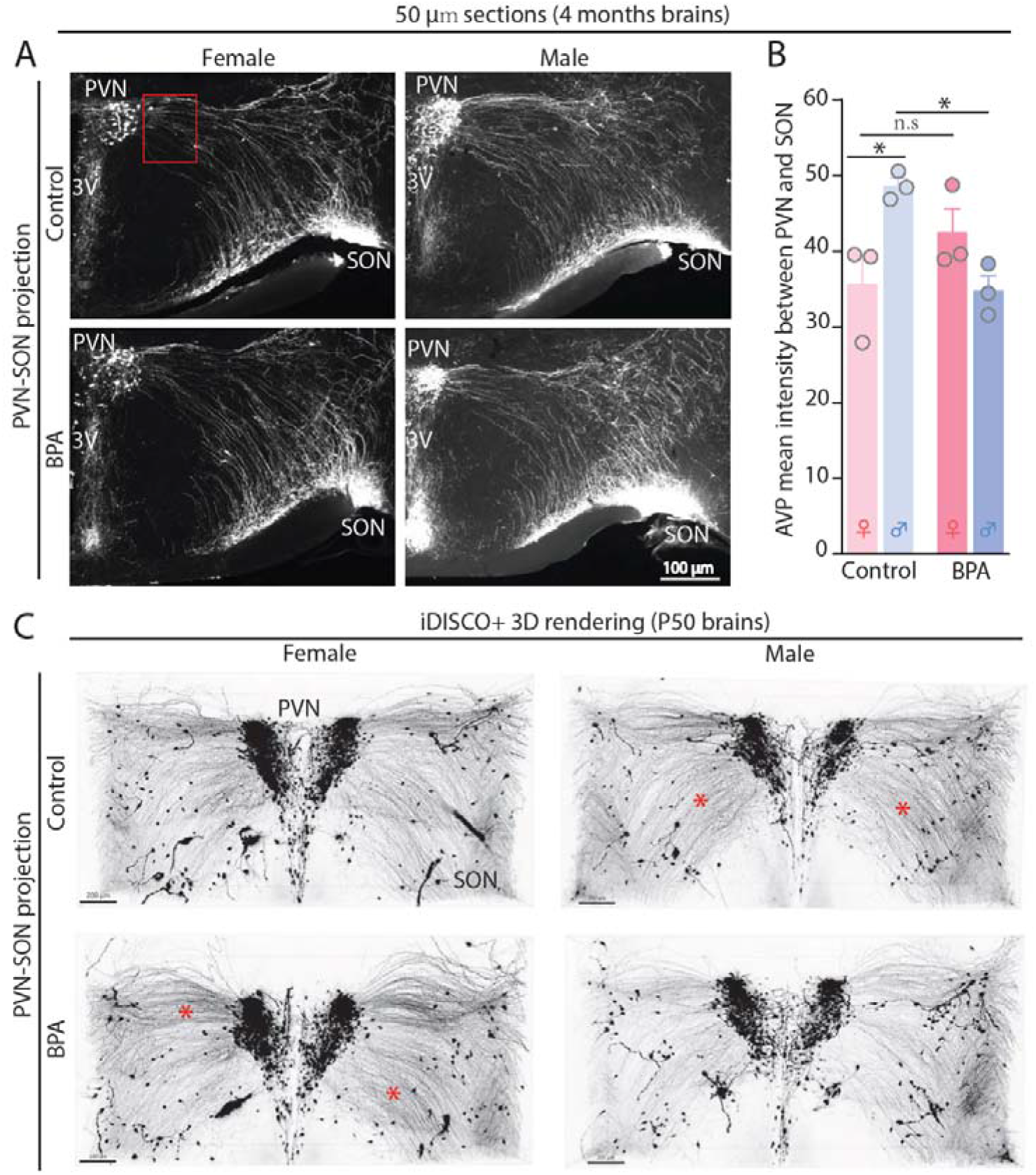
Hypothalamic AVP-ir projections are higher in the adult control male than female and is reversed by gestational exposure to BPA. (**A**) Representative images for AVP intensity between PVN and SON of 4-month female and male brains maternally exposed to corn oil (Control) or BPA. PVN, paraventricular nucleus; SON, supraoptic nucleus; 3V, 3^rd^ ventricle. Red square defines quantification region. (**B**) The quantification of AVP intensity in 50 μm sections of 4-month-old female and male brains. n = 3 mice from 1 litters/sex/group. (**C**) Representative 3D rendered images of AVP projections between PVN and SON presented by iDISCO+ clearing protocol. Asterisks highlight regions of increased AVP intensity compared to control female brains. \**P*< 0.05, n.s. = no significance Data expressed as mean ± SEM, with data analyzed using a one-way ANOVA followed by Sidak’s multiple comparison correction.

Across extra-hypothalamic regions, we observed significantly more AVP-ir in male brains within the lateral septum, BNST, amygdala and lateral habenula (Fig. 5A-H), as expected (*3, 4*). Gestational BPA exposure in males significantly decreased AVP-ir (Fig. 5A-H) and iDISCO AVP+ projections (fig. S2B-C) in the BNST and amygdala but not the lateral septum or lateral habenula (Fig. 5J; fig. S2A). We also observed an increase in amygdala AVP-ir in females exposed to gestational BPA (Fig. 5E and Fig. 5G). Additionally, iDISCO+ revealed other unappreciated extra-hypothalamic regions that display sex differentiation with males having denser AVP projections than the females, including the ventral septal area (VSA) and organum vasculosum laminae terminali (OVLT) in the basal ganglia and ventral forebrain, as well as the paraventricular thalamic nucleus (PV) and mediodorsal thalamic (MD) in the thalamus region (Fig. 5J). Maternal BPA exposure disrupted this dimorphism by increasing the AVP intensity in females and decreasing that in males for the VSA, OVLT and MD (Fig. 5I).

**Fig. 5.**
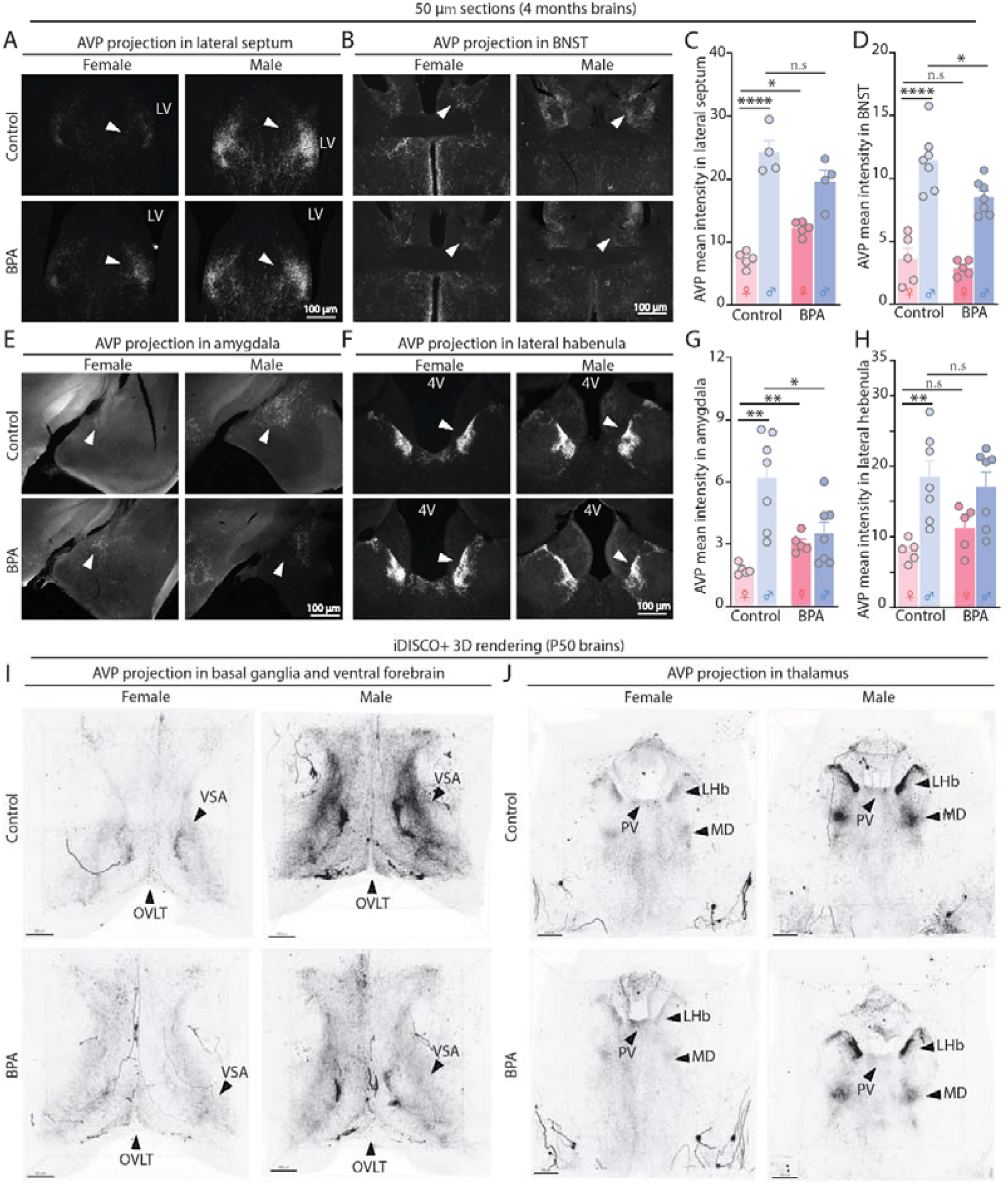
Sexual dimorphism of AVP-ir projections in extra-hypothalamic regions is reversed by gestational exposure to BPA. (**A** and **B**) Representative images of AVP projections in the lateral septum and the BNST in 4-month-old female and male brains maternally exposed to corn oil (Control) or BPA. BNST, the bed nucleus of the stria terminalis; LV, lateral ventricle; arrowheads indicate regions of AVP intensity quantification. (**C** and **D**) Quantitative results of AVP intensity from lateral septum and BNST. n = 4-7 mice from 2 litters/sex/group. (**E** and **F**) Representative images of AVP projections in the amygdala and the lateral habenula of 4-month-old female and male brains maternally exposed to corn oil (Control) or BPA. 4V, 4^th^ ventricle; arrowheads indicate regions of AVP intensity quantification. (**G** and **H**) Quantitative results of AVP intensity from amygdala and lateral habenula. n = 5-7 mice from 2 litters/sex/group. Brown-Forsythe and Welch ANOVA followed by FDR Benjamini and Hochberg correction was used to analyze the amygdala data. Representative 3D rendered images of AVP projections in extrahypothalamic regions the basal ganglia and ventral forebrain (**I**) and thalamus (**J**) of P50 female and male brains exposed to corn oil (Control) or BPA. Arrowheads indicate the specific regions. VSA, ventral septal area; OVLT, organum vasculosum laminae terminali; PV, paraventricular thalamic nucleus; LHb, lateral habenula; MD, mediodorsal thalamic nucleus. *\*P* < 0.05, \*\**P*< 0.01, \*\*\*\**p*< 0.0001, n.s. = no significance; data expressed as mean ± SEM, with data analyzed using a one-way ANOVA followed by Sidak’s multiple comparison correction, unless otherwise stated.

### AVP^PVN^ neurons display sexually dimorphic intrinsic properties that were reversed by maternal BPA exposure

Next, we tested for sexually dimorphic intrinsic and extrinsic electrophysiological properties of AVP^PVN^ neurons. In whole-cell patch-clamp recordings from control animals, male AVP^PVN^ neurons displayed a lower spontaneous action potential (sAP) frequency compared to female AVP+ neurons. Gestational BPA eliminated this sexual dimorphism (Fig. 6A-B). Control male AVP^PVN^ neurons also displayed a more hyperpolarized resting membrane potential than female AVP^PVN^ neurons, indicating females AVP^PVN^ neurons are more excitable than males (Fig. 6C). Maternal BPA exposure had no effects on resting membrane potential in females or males when compared to their same-sex controls (Fig. 6C). We observed no differences in firing rates between control males and females in response to depolarizing current steps (Fig. 6D). Following gestational BPA exposure, female AVP^PVN^ neurons exhibited lower intrinsic excitability than control group (Fig. 6E-F) but no effect was observed in males exposed to prenatal BPA (Fig. 6G); in the control group, male and female AVP^PVN^ neurons displayed similar latencies to first action potential (Fig. 6H), consistent with magnocellular neuronal properties (*21*) however, AVP^PVN^ neurons in female mice gestationally exposed to BPA displayed an longer latency to first spike than males (Fig. 6H-I, and fig. S3A-B). No sexual dimorphism or BPA effects were observed for the Ra, Rm or capacitance measures (Fig. 6K-M). There was no sex difference in the current-voltage and H-current between control males and females; however, female-specific alterations were noted in the gestational BPA exposed groups (fig. S3C-F). These observations reveal a sexual dimorphism between AVP^PVN^ neurons in control animals that is disrupted by BPA. Specifically, female AVP^PVN^ neurons are more excitable than male AVP^PVN^, but this dimorphism is eliminated following gestational BPA.

**Fig. 6.**
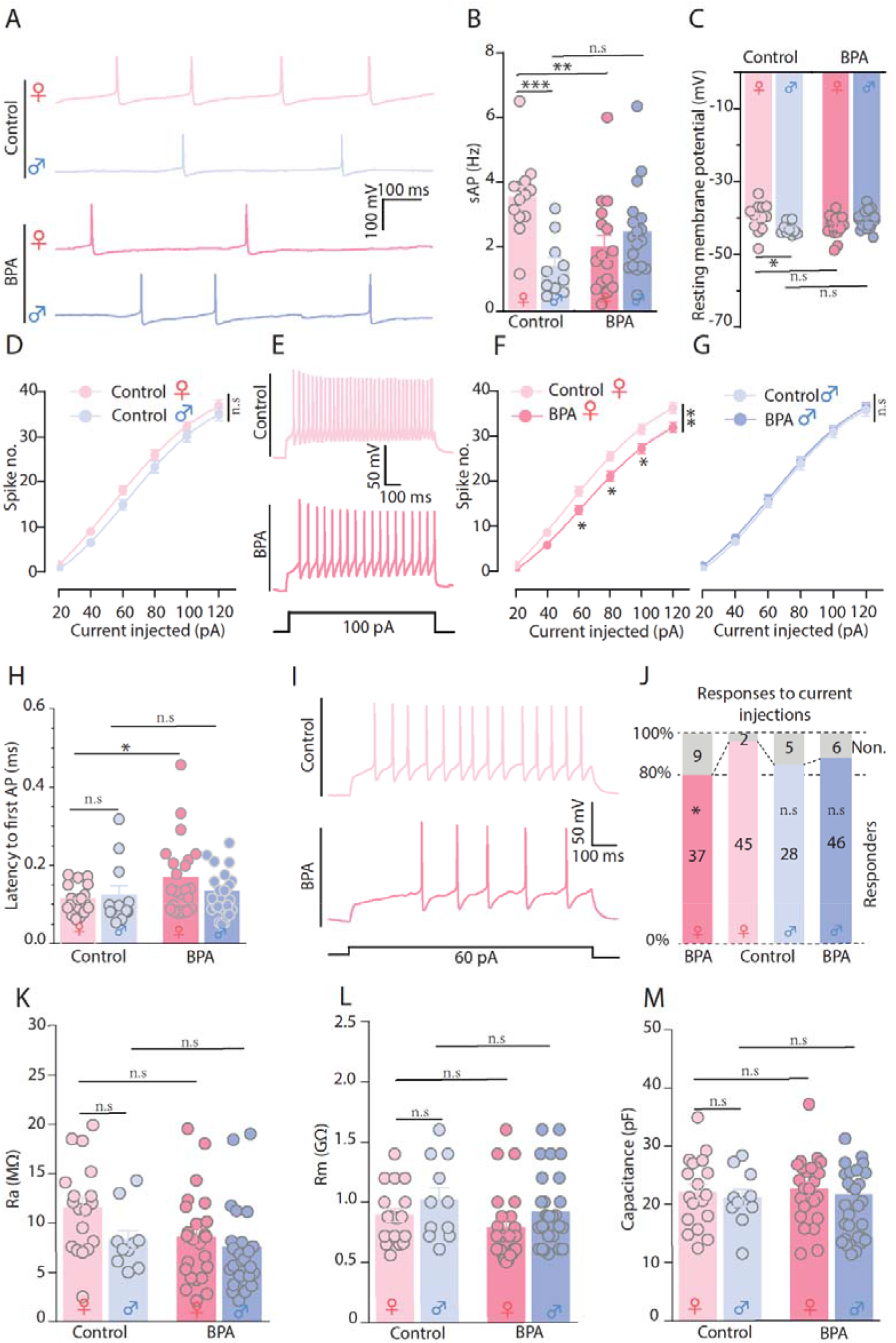
Juvenile AVP^PVN^ neurons display sexually dimorphic intrinsic properties that are disrupted by gestational BPA exposure in females. (**A**) Representative traces and quantification for sAP (**B**) of AVP^PVN^ neurons from female (red) and male (blue) slices in the absence/presence of prenatal BPA exposure. n = 10-16 neurons from 6-7 mice from 4-6 litters/sex/group for sAP. sAP, spontaneous action potential. (**C**) The quantification of resting membrane potential of AVP^PVN^ neurons. n = 10-18 neurons from 6-9 mice from 4-7 litters/sex/group for resting membrane potential. (**E**) Representative firing traces for female AVP^PVN^ neurons in the presence of 100 pA current injection. (**D**, **F** and **G**) Quantitative spike number as a function of current injections. n = 33-52 neurons from 7-11 mice from 4-7 litters/sex/group. Two-way ANOVA repeated measurement analysis followed by Dunnett’s multiple comparison test was used for main group effect, two-way ANOVA repeated measurement analysis followed by Tukey’s multiple comparison test was used for simple effects within timepoints. (**H**) Quantification of the latency to the 1^st^ spike in the presence of 60 pA current injection. n = 12-38 neurons from 4-8 mice from 2-5 litters/sex/group. (**I**) Representative traces for the female AVP^PVN^ neurons showing the latency to the 1^st^ spike evoked by 60 pA current injection (**J**) The percentage of responder AVP^PVN^ neurons under the 40 pA current injection in female and male neurons with/without BPA exposure. n = 33-52 neurons from 7-11 mice from 4-7 litters/sex/group. Gray bars indicate the proportions that did not respond to current injections. Chi-square test was used to test the significance between control females and males, and the BPA effects within same sex. (**K** to **M**) The effects of BPA exposure on the Ra, Rm and capacitance of AVP^PVN^ neurons. n = 10-36 neurons from 4-5 mice from 2-3 litters/sex/group. Ra, access resistance; Rm, membrane resistance. \**P*< 0.05, \*\**P* < 0.01, \*\*\**P*< 0.001, n.s. = no significance; data expressed as mean ± SEM, with all data analyzed using one-way ANOVA followed by Sidak’s multiple comparison correction, unless otherwise stated.

We also assessed synaptic transmission onto male vs female AVP^PVN^ neurons and observed no significant sex difference in spontaneous excitatory postsynaptic potential (sEPSC) of control animals. Gestational BPA exposure increased the frequency but not amplitude of sEPSCs in female AVP^PVN^ neurons (fig. S4A-D). No sex differences were observed in spontaneous inhibitory postsynaptic potential (sIPSC) between male and female AVP^PVN^ neurons +/- gestational BPA (fig. S4E-G). There was no difference in the sEPSC/sIPSC ratio (fig. S4H-I).

## Discussion

Our study demonstrates that hypothalamic AVP sex differentiation occurs from neuronal birth, with two-thirds more AVP+ neurons born at E11 in male than female PVN. Consistent with rodent studies in the hippocampus whereby endogenous estrogen promote sex-biased neonatal neurogenesis (*22, 23*), we show estrogen is necessary and sufficient to induce AVP^PVN^ sex differentiation in the mouse embryonic hypothalamus. Debate continues as to whether estrogen or testosterone is the masculinizing signal in the human brain; although a recent study shows that testosterone promotes the proliferation of radial glial progenitors in a sex-specific manner in human brain organoids, whereas estrogen drove sex differentiation in mice brain organoids (*24*). Additional studies are needed into species variations in steroid hormones underlying sex-specific brain development as well as to determine the importance of hormone-driven neurogenesis as a mechanism driving sex differentiation in other brain regions.

In mice, current dogma states that masculinization is driven by a perinatal surge of gonadal testosterone that is converted to estradiol in sexually dimorphic brain regions (*25*). We show that restricting gestational BPA exposure to the neurogenic window (E7.5-E15.5) causes females to display male-like AVP+ numbers in adolescence (P15), suggesting that an embryonic estrogen cue has permanent effects. This result raises intriguing questions as to the interplay between the sexually dimorphic organizing influences of embryonic versus perinatal hormone induction signals. Moreover, in humans, neural sex differentiation is thought to occur during the second trimester, when testosterone levels are higher in male than female fetuses (*26*). Coincidentally, neurogenesis is the core neurodevelopmental process during the second trimester (*27*). Our findings here uncover the possibility that mice and humans utilize a similar neurogenic mechanism for fetal brain sex differentiation, opening new lines of research in mice to study human programs.

These findings raise unanswered questions regarding from where this estrogenic inductive cue in male mice arises given that the testes only start to differentiate from the genital ridge at E12 and secrete non-physiological levels (<1 pmole/testis) of intratesticular testosterone for further testis development until near birth (>E17), when plasma levels rise (*28*). High levels of maternal estrogens can enter fetal circulation; however, this is an unlikely source since alpha fetoprotein binds estrogen in fetal circulation and sequesters it from entering the brain (*29*). Instead, de novo neural sex steroid synthesis from membrane-bound cholesterol and/or sex chromosome genes are candidate mechanisms. Evidence now shows that the adult brain is a steroidogenic site independent from the gonads, with neurosteroidogenesis evolutionary conserved across fish, birds, and mammals (*30, 31*). Despite fetal neurosteroidogenesis proposed to contribute to differential neurodevelopment (*32*), its role remains entirely unexplored. Moreover, the intrinsic expression of sex chromosome genes have underappreciated differential cellular effects (*33*). Although less recognized, Y- encoded genes expressed in the brain can have male-specific effects (*34, 35*), whereas XX cells contain a paternally imprinted X that can contribute to cell-autonomous differences in female cells (*36*). Interestingly, AVP-ir is higher in *XY^-sry^* males that develop ovaries than XX^*+sry*^ males with testis or intact XX females, suggesting genes on the Y chromosome might influence vasopressin sexual dimorphisms aside from sex hormones.

Despite agreement that vasopressin is sexually dimorphic across several regions that control complex social behaviors (*3, 4*); evidence is incomplete as to whether AVP is dimorphic in the hypothalamic PVN, which is the predominant producer of this neuropeptide in the brain (*3*). Here we show convincingly that the PVN is sexually dimorphic for AVP+ cell bodies, projections, and intrinsic electrophysiological properties, and that embryonic exposure to the xenoestrogen BPA permanently disrupts the sexually dimorphic development of this hypothalamic AVP system in adult offspring. These data also point out, once again, the adverse effects of environmentally relevant levels of BPA on developing brains and bring new insights into the mechanism by which maternal BPA exposures might associate with neurodevelopmental and psychiatric disorders in children.

## Acknowledgements

We thank the Kurrasch Lab for input during this study and appreciate Natalia Klenin’s help with cryo-sectioning and Dr. David Elliott’s assistance with light-sheet imaging in the HBI AMP at University of Calgary.

## Funding

This work was supported by fundings listed below:

Canadian Institutes of Health Research grant 470608 (DMK)

Canadian Institutes of Health Research grant FDN-148440 (JB)

Natural Sciences and Engineering Research Council of Canada grant DG386445 (DK)

New Frontiers in Research Fund NFRF-2018-00597 (CL and DMK)

University of Calgary Owerko Centre Postdoctoral Fellowship (JZ)

Natural Sciences and Engineering Research Council BRAIN CREATE Postdoctoral

Fellowship (JZ)

Canadian Institutes of Health Research Postdoctoral Fellowship MFE-183073 (JZ)

## Author contributions

Conceptualization: JZ, DK

Methodology: JZ, PM, DB, JB, DK

Investigation: JZ, PM, DB

Visualization: JZ, PM

Funding acquisition: CL, JB, DK

Project administration: DK

Supervision: DK

Writing – original draft: JZ, DK

Writing – review & editing: JZ, PM, DB, CL, JB, DK

All the authors contributed to intellectual discussion and the direction of the project.

## Data and materials availability

All data are available in the main text or the supplementary materials.

## Supplementary Materials

Materials and Methods

Figs. S1 to S4

References (37-40)

## Supplementary Materials for

### Materials and Methods

#### Animals

All animal procedures and protocols were approved by the University of Calgary Animal Care and Use Committee. All the experiments, including the development of AVP neurons, intra-/extra-hypothalamic AVP projections, ÌDISCO+ tissue clearing, neurogenesis, and *in utero* injections were conducted in CD1 male and female mice, except for whole-cell patch clamp studies, which were conducted in the offspring obtained by crossing AVP-IRES2-Cre-D (JAX #023530) with Rosa26^tdTomato^ (JAX #007914). CD1 and C57 female and male mice aged 6-8 weeks were obtained from Charles River Laboratories and the Jackson Laboratory (JAX), respectively, and were housed on a 12-h:12-h light: dark cycle (lights on at 6:00 a.m.). Animals were acclimatized for seven days before being bred. Embryonic day 0.5 (E0.5) was recorded on the day a vaginal plug was observed. For animals receiving BPA, the standard diet (4.6 kcal/g) was replaced with BPA (2.25 μg/kg bw/day) (TD. 200755, Envigo) or 7% corn oil laced diet (TD. 120176, Envigo) on days described in the results.

#### Genotyping

Tail samples of the mice were collected, incubated with 500 μl of tail buffer containing proteinase K (1:200, p8107S, New England Biolabs) at 37 □ for overnight to lyse the cellular structure. Equal volume of isopropanol (I9516, Sigma) was added to the lysate and the resultant was then centrifuged at 13,200 rpm for 10 min at 4□. After a 70% ethanol wash of the genomic DNA, samples were centrifuged at max speed for 10 min at 4□. DNA was then dissolved in ddH_2_O until used for sex-determining region Y (SRY) genotyping to distinguish/confirm the sexes of the animals. As previously described(37), *OneTaq^®^* Master Mix (M0486L, New England Biolabs) was used to carry out the polymerase chain reaction (PCR) according to the manufacturer’s instructions, with the following primers: Sry, 5’- TCATGAGACTGCCAACCACA-3’ and 5’-CATGACCACCACCACCACCA-3’. Amplified DNA products were electrophoresed in the 2% agarose gel and imaged with a Gel Doc XR+ system (Bio-Rad). Each sample were run for two to three times to ensure the accuracy of sex determination.

#### Neurogenesis assay

Pregnant mice exposed to BPA or corn oil were injected with the thymidine analog 5-bromo-2’-deoxyuridine (BrdU, 100 mg/kg, B5002, Sigma) at E10.5, 11.5 and 12.5 to label the cells born on the day of injection. The brains were collected at P0, and the procedures of tissue preparation and Immunohistochemistry follows those described below.

#### Tissue preparation

Prior to collecting the brains, animals (postnatal day 15 (P15), P30 and 4 months) were subjected to cardiac perfusion with ice-cold PBS and 4% paraformaldehyde (PFA) consecutively. The brains were then immersed in 4% PFA overnight at 4 □ followed by three washes with PBS and equilibration in 20% sucrose/PBS overnight at 4 □. The brains collected at E15.5, P0 and P7 were directly fixed with 4% PFA overnight at 4 □. After being embedded in OCT (CA95057-838, VWR), the brains were either coronally sectioned (10 μm sections) on a cryostat (CM3050 S, Leica) or sliced (50 μm sections) on a vibratome (VT1200S, Leica). The 10 μm sections were collected on Superfrost plus glass micro-slides (12-550-15, Fisher Scientific). Serial sectioned tissue containing the hypothalamus was collected over 10-12 slides (10 slides for E15.5, 12 slides for other time points). While the 50 μm sections were collected in the 48-well plates, every-other sections from the same brain were stained for specific antibodies.

#### Immunohistochemistry

For the 10 μm sections, tissue was rehydrated in PBS for 10 min, washed for 3 x 10 min with PBT (0.1% Triton X-100 in PBS), permeabilized with 1% Triton X-100 in PBS for 30 min, blocked with 5% normal donkey serum (NDS, D9663, Sigma) in PBT for 1 hr at room temperature, and incubated with rabbit anti-vasopressin (1: 500, ab213708, abcam) and sheep anti-BrdU (1:500, ab1893, abcam) in PBST with 5% NDS overnight at 4□. The slides were then washed with PBT for 3 x 10 min at RT, followed by incubating with secondary antibodies (1: 500, Alexa 488 donkey anti-rabbit, A21206; Alexa 555 donkey anti-sheep, A21436) in PBT with 5% NDS for 1 hr at RT. Slides were then washed with PBT for 3 x 10 min at RT, followed by staining with Hoechst 33342 (1:1000, H3570, Life Technologies) in PBS for 10 min at RT. The stained slides were then mounted with Aqua Poly/Mount (18606, Polysciences). For the birthdating experiments, an additional DNA hydrolysis step (slides were treated with 2N HCl for 30 min at 37 □) followed by permeabilization to allow the anti-BrdU antibody access to the BrdU within the DNA was conducted. For the projection experiments, 50 μm sections were stained using a free-floating approach. All the procedures were the same as the slide mounted protocol, except for the durations of the permeabilization, blocking, primary antibody and secondary antibody incubation, which were 45 min, 1.5, 16 and 2.5 hr, respectively. In addition, the anti-AVP primary antibody dilution was 1:1000 for the free-floating protocol.

#### Imaging

Images of immunostained sections were captured using a ZEISS Axioplan2 manual compound microscope (10X or 20X objectives) equipped with a Lumen Dynamics X-Cite Series 120 Q fluorescent module and ZEISS Axiocam HRc camera or a ZEISS LSM 880 with Airyscan confocal microscope (20X objective). Images were preprocessed using the Zen Black software. Image J software (version 1.49s; U.S. National Institutes of Health) was used to adjust the brightness and/or contrast of the entire image if appropriate.

#### iDISCO+ tissue clearing, staining, and imaging

The iDISCO+ protocol was described previously(38). Briefly, P7 and adult CD1 mice were anesthetized and cardiac perfused with PBS and 4% PFA before harvesting brains. The brains were then immersed in 4% PFA overnight with shaking at 4 □, followed by 1 hr at RT. The brains were washed with PBS for 3 x 30 min with shaking at RT prior to dehydration with methanol/H_2_O series: 20%, 40%, 60%, 80%, 100%, 100% for 1 hr each at RT, followed by overnight incubation in 66% DCM (Dichloromethane, 270997, Sigma)/33% methanol (A412SK-4, Fisher) with shaking at RT. The brains were then washed with 100% methanol for 2 x 1 hr at RT, chilled for 15 min at 4 □, and bleached with 5% H_2_O_2_/methanal overnight at 4 □. The samples were rehydrated with a methanol/H_2_O series: 80%, 60%, 40%, 20%, PBS for 1 hr each at RT, and washed in PTx.2 (2% triton X-100 in PBS) for 2 x 1 hr at RT before utilized for immunostaining. The immunolabeling step consists of permeabilizing in permeabilization buffer (20% (v/v) DMSO and 2.3% (w/v) glycine in PTx.2) for 2 days at 37 □, blocking in the blocking buffer (10% (v/v) DMSO and 6% NDS in PTx.2) for 2 days at 37 □, incubating with rabbit anti-vasopressin (1:400, ab213708, abcam; 13 days for P7 brains, 33 days for P50 brains) in PTwH (0.2% Tween-20 and 10 μg/ml heparin in PBS)/5%DMSO/3% NDS at 37 0, washing in PTwH for 5 x 1 hr at 37 0, incubating with secondary antibody Alexa 555 donkey anti-rabbit (1:200, ab31572, abcam; 13 days for P7 brains and 27 days for P50 brains) in PTwH/3% NDS at 37 □, and washing in PTwH for 5 x 1 hr at 37 □. The brains were then dehydrated in methanol/H_2_O series: 20%, 40%, 60%, 80%, 100%, 100% for 1 hr each at RT, incubated with 66% DCM/33% methanol with shaking for 3 hr at RT, incubated with 100% DCM for 2 x 15 min at RT to wash the methanol, added to ethyl cinnamate (ECi) with shaking for over 2 hr at RT. The cleared brains were then imaged with a LaVision BioTec UltraMicroscope II light-sheet microscope equipped with Olympus MVPLAPO 2x/0.50 NA with 5.7, 6 and 10 mm WD dipping caps and Andor Zyla 4.2 Plus sCMOS camera, with a step size of 3 μm and 20 μm. The images were processed and analyzed with LaVision BioTec ImSpector (Miltenyi Biotec) and Imaris (version 9.9.0).

#### *Ex Vivo* electrophysiology

*Slice preparation*. AVP-IRES2-Cre-D; Rosa26^tdTomato^ male and female mice maternally exposed to BPA or corn oil were anesthetized and decapitated at the age of P30 to P40. Brains were rapidly removed and placed in the ice-cold slicing solution containing (in mM) 87 NaCI, 2.5 KCI, 0.5 CaCl_2_, 7 MgCl_2_, 25 NaHCO_3_, 25 D-glucose, 1.25 NaH_2_PO4 and 75 sucrose bubbled with 95% O_2_/5% CO_2_. Vibratome (VT1200S, Leica) was used to obtain coronal sections that contain PVN. Obtained sections were allowed to recover incubating for 1 hr in prewarmed (30 □) artificial cerebrospinal fluid (aCSF) containing (in mM) 126 NaCl, 2.5 KCl, 26 NaHCO_3_, 2.5 CaCl_2_, 1.5 MgCl_2_, 1.25 NaH_2_PO4 and 10 glucoses, bubbled with 95% O_2_/5% CO_2_. *Electrophysiology*. All the recordings were carried out in aCSF at 30-32 □ perfused at a flow rate of 1 mL/min. Neurons were visualized using an upright microscope fitted with differential interference contrast and epifluorescence optics (UVICO, RappOptoElectronics) and camera (AxioCam MRm). Borosilicate pipettes (3-5 mΩ) were filled with the internal solution containing (in mM): 108 K-gluconate, 2 MgCl_2_, 8 Na-gluconate, 8 KCl, 1 K_2_-EGTA, 4 K_2_-ATP, 0.3 Na_3_-GTP, 10 mM HEPES, and whole-cell patch-clamp recordings were made using an Axopatch 200B amplifier (Molecular Devices) and pClamp 10.2 (Molecular Devices). Prior to any recording, the neurons were allowed to stabilize at −60 mV in voltage clamp mode for 2 min. In the current clamp recordings, spontaneous action potentials were recorded at resting membrane potential. All the current steps protocols were conducted at the initial membrane potential of −70 mV, except the H-current protocol which was with an initial membrane potential of −50 mV. In the voltage clamp recordings, the spontaneous excitatory postsynaptic currents (sEPSC) and spontaneous inhibitory postsynaptic currents (sIPSC) were recorded when the neurons were held at −70 mV and 0 mV, respectively. Access resistance (Ra) was consistently monitored, and recordings were used for analysis only if Ra changes were < 15‰

#### 17β-estradiol and fulvestrant injection

The injections of 17β-estradiol (4 μM; E8875, Sigma) and fulvestrant (4 μM, I4409, Sigma) were adapted from the *in utero* electroporation procedures described elsewhere(*39, 40*). Briefly, pregnant CD1 mice exposed to BPA or corn oil were anesthetized with 5mL/min of isoflurane (CP0406V2, Fresenius Kabi), followed by 2.5 mL/min to maintain the anesthesia during surgery, with the oxygen flow rate of 1 mL/min. Baytril (DIN 02169428, Bayer) and buprenorphine (DIN 02342510, Ceva Animal Health Inc.) were administered before the surgery. Approximately 2 μL of 17β-estradiol and fulvestrant mixed with fast green (FG, 0.05 mg/mL in aCSF, F7252, Sigma) were injected into the lateral ventricles of the E13.5 brains with an Eppendorf FemtoJet 4i microinjector (VWR) and a Narishige 3-axis M152 micromanipulator (Leica). The injected embryonic brains were gently massaged with the forceps until the visualization of FG in the third and fourth ventricles. The uterus containing embryos were then placed back inside the abdominal cavity followed by filling it with prewarmed sterile saline. The abdominal walls and the skin were sutured before the anesthesia was stopped. The dams were then subcutaneously injected with Ringer’s solution (2 mL) and placed on a heat pad to allow prompt recovery. The injected embryonic brains were collected at E15.5 to conduct immunochemical experiments as per the protocol described above.

#### Quantification and statistics

For cell count, 10 μm of coronal sections from one of ten (E15.5 brains) or one of twelve slides (postnatal brains) were stained and quantified manually using the Image J (version 1.49s; U.S. National Institutes of Health) and the summation of specific neuronal type number were used for analysis. For AVP projections, 50 μm sections were collected and stained for every other section, three sections of ROI were used for intensity quantification, and the summation of mean intensity of each section were used for analysis. All the data were displayed as mean ± SEM, and unpaired t-test (two-tailed), or ordinary one-way ANOVA or repeat measurement two-way ANOVA were used to analyze the difference between groups using GraphPad Prism 8.02 (GraphPad Software). The statistical significance threshold is 0.05.

**Fig. S1.**
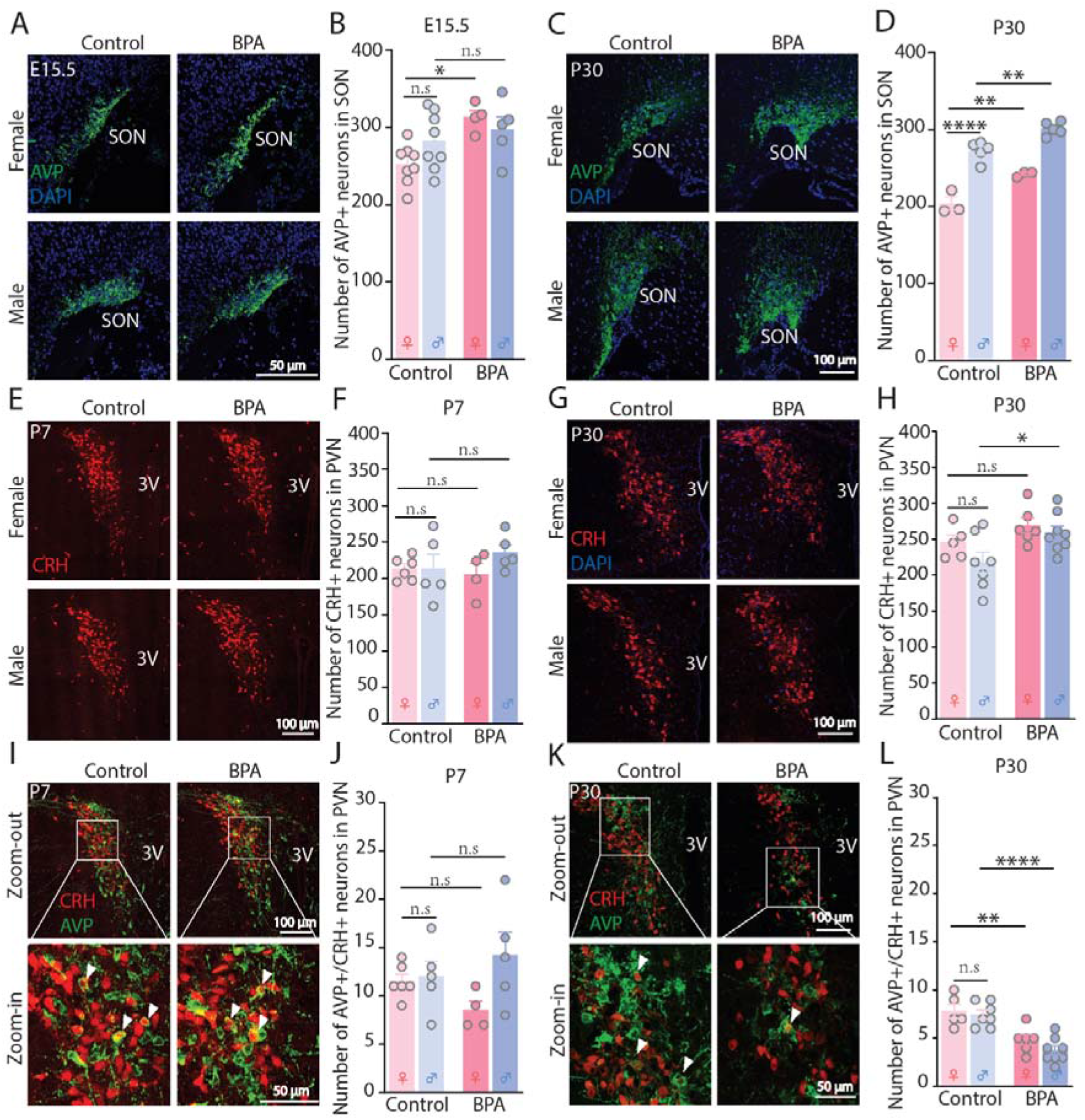
The effect of gestational BPA exposure on AVP+ neurons in other hypothalamic nuclei or other PVN cell types. (**A** to **D**) The effect of BPA exposure on the number of AVP neuron in the E15.5 and P30 female and male SON. n = 3-8 mice from 1-2 litters/sex/group. SON, supraoptic nucleus. (**E** to **H**) The effect of BPA exposure on the number of CRH neuron in the P7 and P30 female and male PVN. n = 5-9 mice from 2-3 litters/sex/group. PVN, paraventricular nucleus; CRH, corticotrophin-releasing hormone; 3V, 3^rd^ ventricle. (**I** to **L**) The impact of maternal BPA exposure on the number of AVP+/CRH+ neurons in the PVN. n = 5-8 mice from 2-3 litters/sex/group. Arrows show the AVP+/CRH+ neurons. \**P* < 0.05, \*\**P* < 0.01, \*\*\*\**P* < 0.0001, n.s indicates no significance; graphs show data expressed as mean ± SEM, and all data were analyzed with one-way ANOVA followed by Sidak’s multiple comparison correction.

**Fig. S2.**
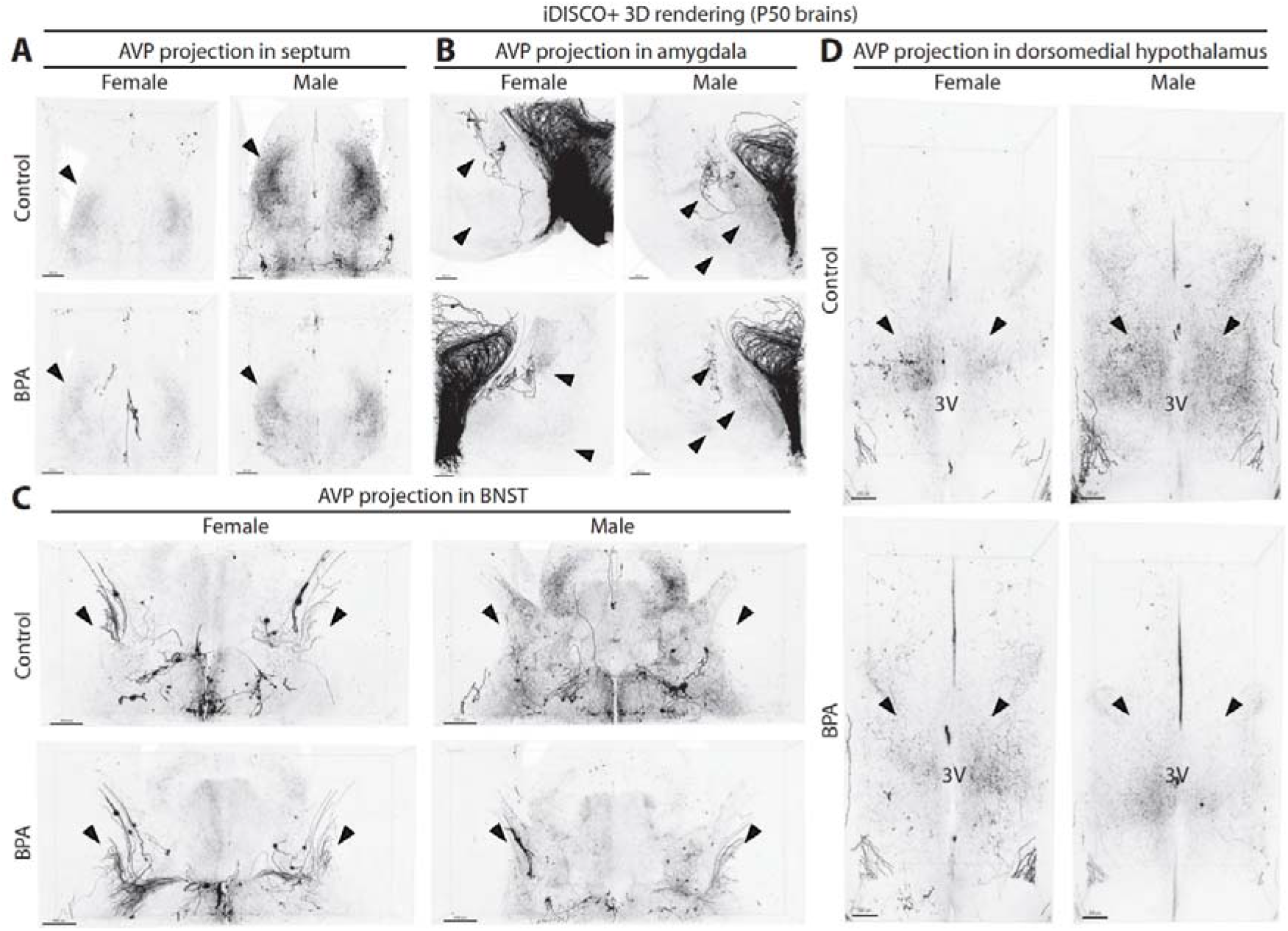
Sexual dimorphic of AVP-ir projections in adult brains is reversed by gestational BPA exposure in cleared whole brains. (**A** to **D**) The representative 3D rendering of AVP projections in the female and male septum, amygdala, BNST and dorsomedial hypothalamus in the absence (Control) and presence of BPA exposure. Arrows show the regions of interest. 3V, 3^rd^ ventricle.

**Fig. S3.**
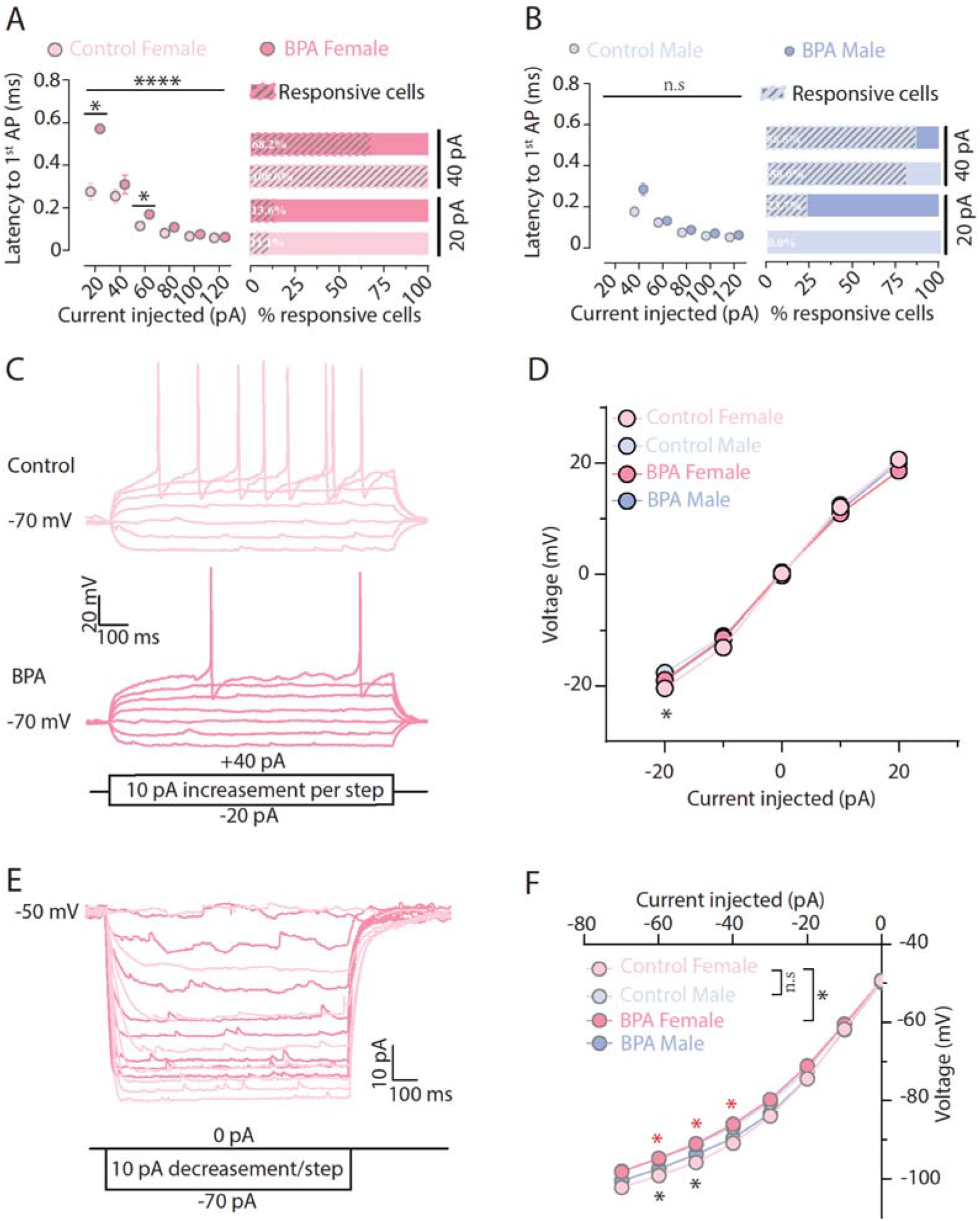
Maternal BPA exposure alters the intrinsic properties of females AVP^PVN^ neurons. Effects of BPA exposure on the latency to the first spike of AVP^PVN^ neurons in female (**A**) and male (**B**) brains in response to current injections. n = 12-38 neurons from 4-8 mice from 2-5 litters/sex/group. These data were analyzed using Mixed- effect analysis. Slash shadow bars indicate the percentage of responsive neurons. (**C**) Representative traces showing the current-voltage relationship of female AVP^PVN^ neurons in the absence/presence of gestational BPA treatment. (**D**) The quantitative results of the effect of BPA exposure on the current-voltage relationship of AVP^PVN^ neurons. n = 11-35 neurons from 4-5 mice from 2-5 litters/sex/group. *P* values in black mean the comparison between control female and control male. Two-way ANOVA repeated measurement analysis followed by Dunnett’s multiple comparison test was used for main group effect, two-way ANOVA repeated measurement analysis followed by Tukey’s multiple comparison test was used for simple effects within injections. (**E**) Representative traces showing the H current of female AVP^PVN^ neurons in the absence/presence of gestational BPA treatment. (**F**) The quantitative results of the effect of BPA exposure on the H current of AVP^PVN^ neurons. n = 12-31 neurons from 4-8 mice from 2-5 litters/sex/group. *P* values in black mean the comparison between control female and control male, while the values in red mean that between control female and BPA female. \**P* < 0.05, \*\*\*\**p* < 0.0001, n.s indicates no significance; graphs show data expressed as mean ± SEM. Two-way ANOVA repeated measurement analysis followed by Dunnett’s multiple comparison test was used for main group effect, two-way ANOVA repeated measurement analysis followed by Tukey’s multiple comparison test was used for simple effects within injections.

**Fig. S4.**
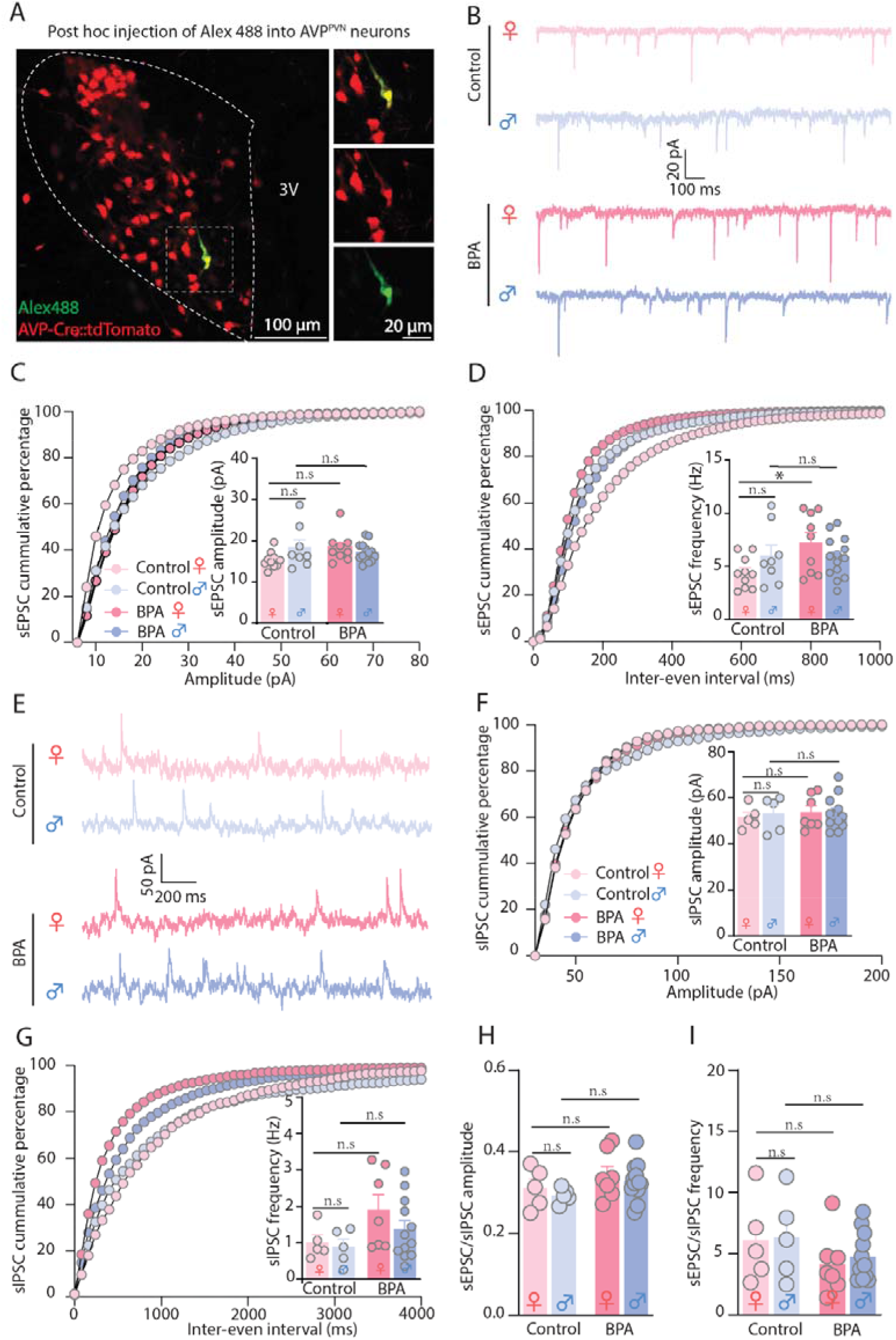
Maternal BPA exposure specifically disrupted the excitatory synaptic transmission of female juvenile AVP^PVN^ neurons. **(A)** Representative images showing *post hoc* injection of Alex 488 into a patch clamped AVP^PVN^ neuron in the PVN slice. **(B)** The representative traces of spontaneous excitatory postsynaptic current (sEPSC) recorded from male and female AVP^PVN^ neurons in the absence/presence of maternal BPA exposure. Impacts of BPA exposure on the amplitude (**C**) and frequency (**D**) of sEPSC. n = 8-14 neurons from 4-5 mice from 2-3 litters/sex/group. (**E**) The representative traces of spontaneous inhibitory postsynaptic current (sIPSC) recorded from male and female AVP^PVN^ neurons in the absence/presence of maternal BPA exposure. Impacts of BPA exposure on the amplitude (**F**) and frequency (**G**) of sIPSC. n = 4-12 neurons from 3-4 mice from 2-3 litters/sex/group. (**H** and **I**) The ratio of sEPSC to sIPSC for amplitude and frequency. n = 4-12 neurons from 3-4 mice from 2-3 litters/sex/group. \**P* < 0.05, n.s indicates no significance; graphs show data expressed as mean ± SEM, and all data were analyzed with one-way ANOVA followed by Sidak’s multiple comparison correction.

